## Supplemental information for "Vangl2 suppresses NF-κB signaling and ameliorates sepsis by targeting p65 for NDP52-mediated autophagic degradation"

**This PDF file includes:**

Figures S1 to S6

Materials and Methods

Supplementary Figures

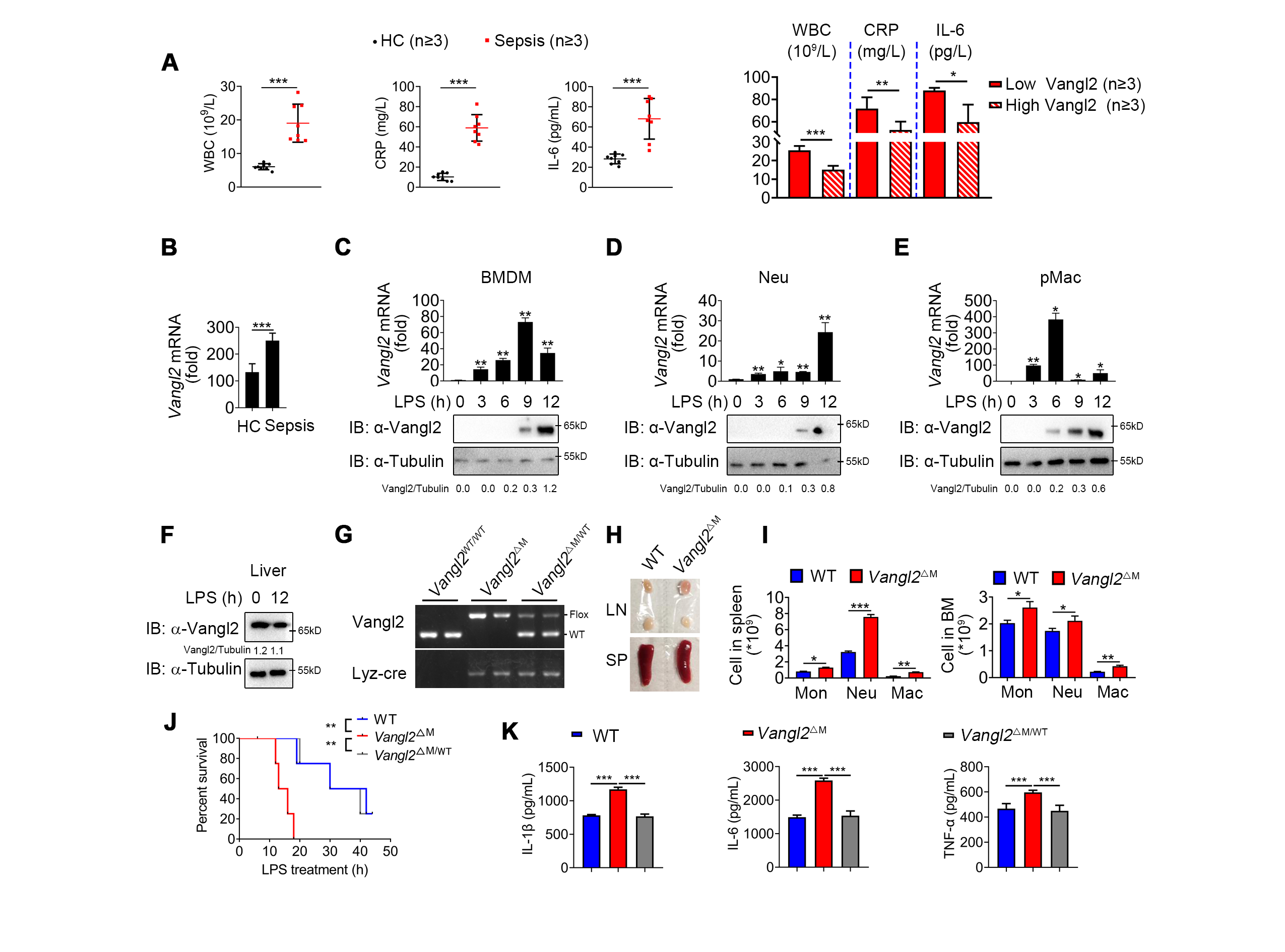

**Figure 1-figure supplement 1. Expression of Vangl2 during sepsis and LPS treatment.**

(A) White blood cell (WBC) count, acute C-reactive protein (CRP), and IL-6 level in the PBMC of healthy volunteers (HC) and sepsis patients (n≥3). And WBC count, CPR, and IL-6 level in the PBMC of low Vangl2 group and high Vangl2 group (n≥3).

(B) Vangl2 expression in spleen of healthy volunteers and sepsis patients was analyzed by GEO (GSE69063, GSE145227, and GSE46955) analysis (n≥5).

(C-E) The mRNA and protein levels of Vangl2 in BMDMs, Neu, or pMac from WT mice after LPS treatment for the indicated times were detected.

(F) The protein levels of Vangl2 in the liver of WT mice after LPS treatment for the indicated times were detected.

(G) Genotyping of transgenic mice by PCR and agarose gel electrophoresis.

(H) Gross images of lymph nodes (LN) and spleens (SP) of WT and *Vangl2^ΔM^* mice.

(I) Flow cytometry analysis of myeloid cell populations in the spleens of WT and *Vangl2^ΔM^* mice (n≥4).

(J) The survival rates of WT, *Vangl2^ΔM^* and *Vangl2^ΔM/WT^* mice treated with high-dosage of LPS (30 mg/kg, i.p.) (n≥4).

(K) IL-6 and TNF-α secretion by WT, *Vangl2^ΔM^* and *Vangl2^ΔM/WT^* BMDMs treated with LPS for 6 h was measured by ELISA. IL-1β secretion by WT, *Vangl2^ΔM^* and *Vangl2^ΔM/WT^* BMDMs treated with LPS for 6 h and ATP for 30 min was measured by ELISA.

BMDM, bone marrow-derived macrophage; Neu, neutrophil; pMac, peritoneal macrophage; Mon, monocyte. Data are representative of three independent experiments and are plotted as the mean ±SD. **p*<0.05, ***p*<0.01, ****p*<0.001 vs. corresponding control.

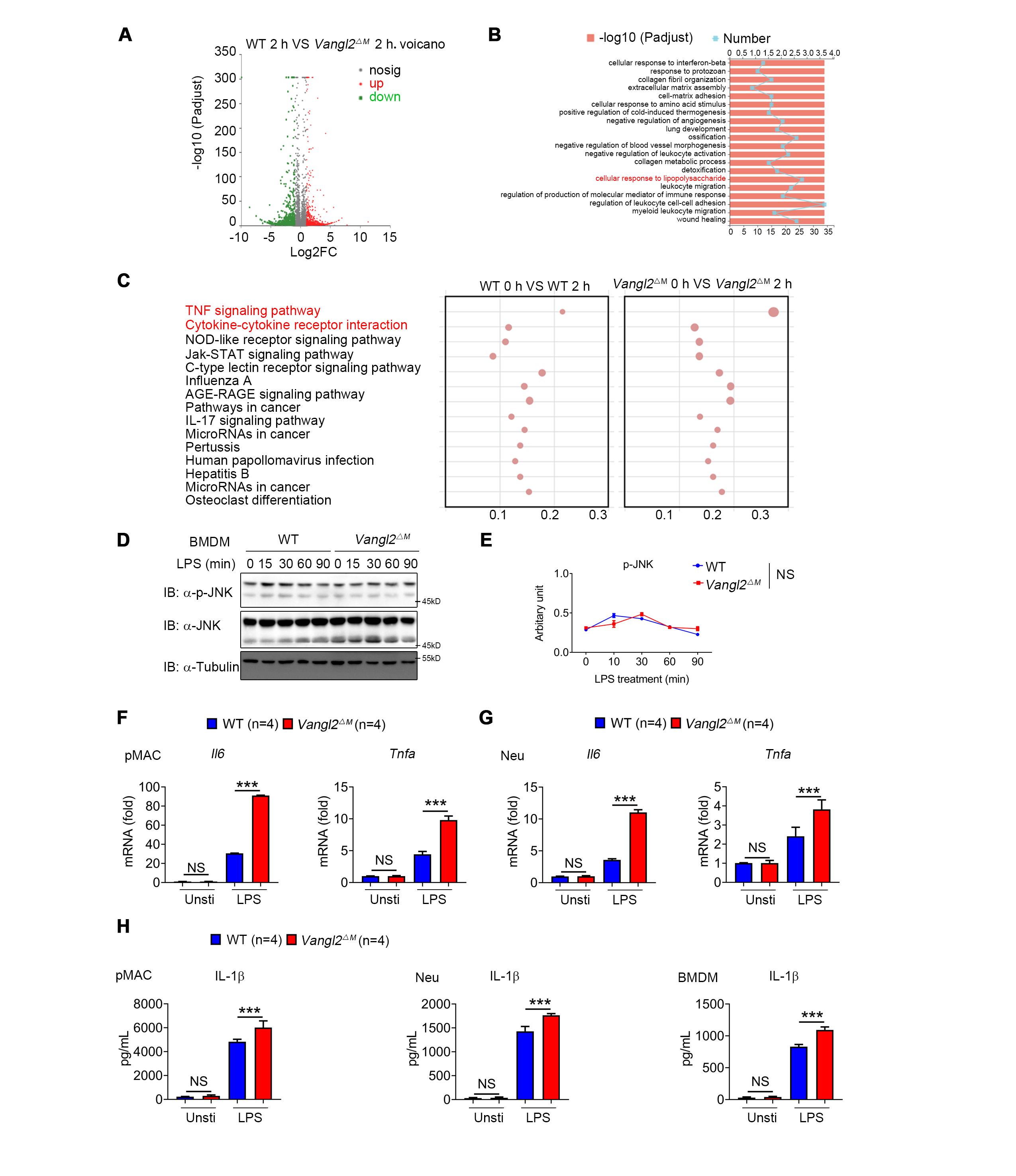

**Figure 2-figure supplement 2. Vangl2 defection promotes LPS-induced NF-κB activation and production of inflammatory cytokines.**

(A-C) Volcano plot (A), GO analysis (B) and KEGG analysis (C) of differentially expressed genes in WT and Vangl2-deficient BMDMs after LPS treatment for 2 h.

(D and E) WT and Vangl2-deficient (n≥3) BMDMs were stimulated with LPS (100 ng/ml) for the indicated times. Immunoblot analysis of total and phosphorylated JNK.

(F and G) WT and Vangl2-deficient peritoneal macrophages and neutrophils were stimulated with LPS for 6 h. mRNA levels of *Il6* and *Tnfa* were measured by qPCR (n=4).

(H) ELISA analysis of IL-1β in the supernatants of WT and Vangl2-deficient BMDMs, neutrophils, or peritoneal macrophages stimulated with LPS for 6 h and ATP for another 0.5 h.

Data are representative of three independent experiments and are plotted as the mean ±SD. **p*<0.05, ***p*<0.01, ****p*<0.001 vs. corresponding control.

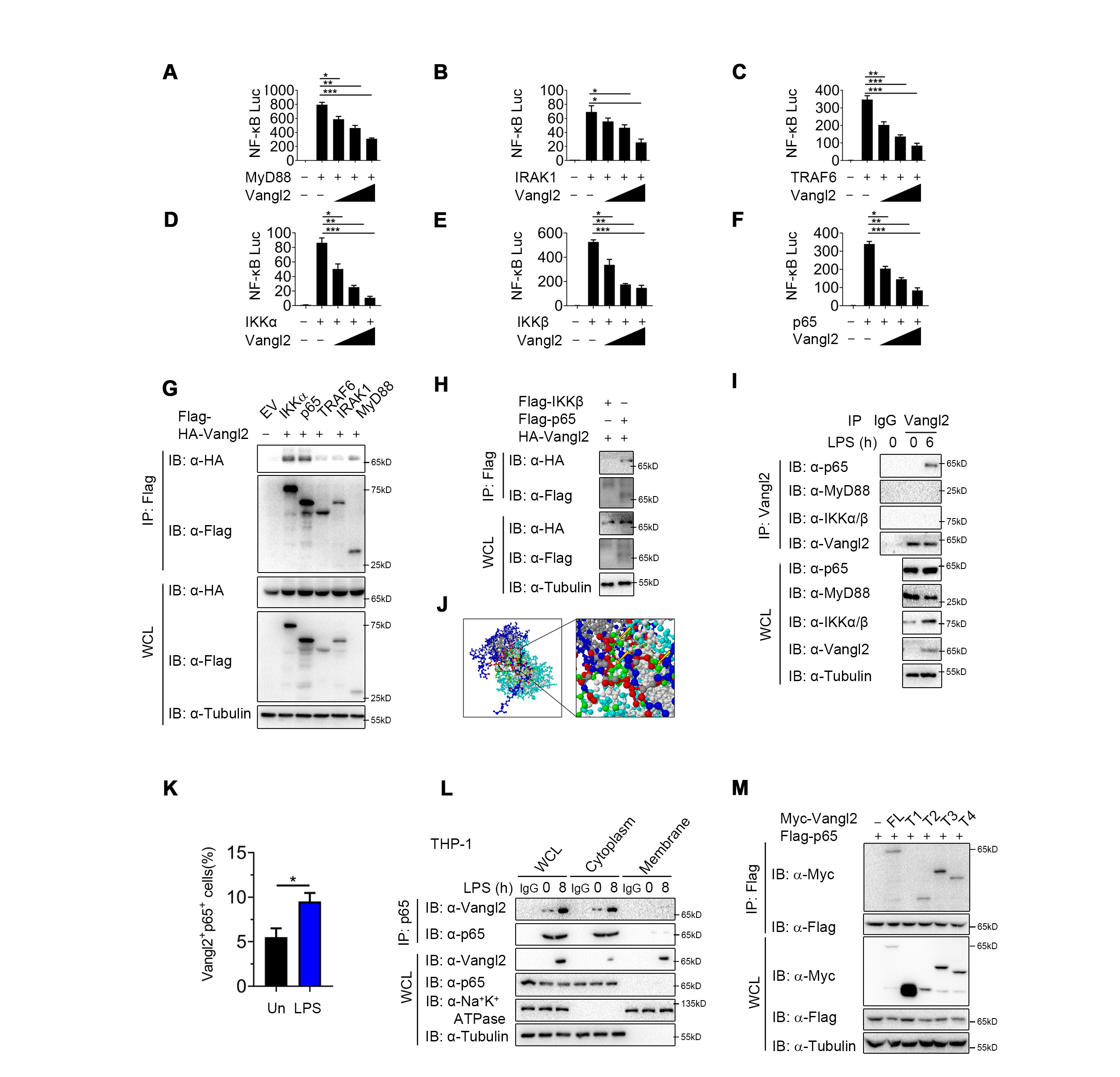

**Figure 3-figure supplement 3. Vangl2 interacts with p65 to inhibit NF-κB activation.**

(A-F) Luciferase activity in HEK293T transfected with plasmids encoding an NF-κB luciferase reporter and TK-Renilla reporter, together with a vector encoding MyD88 (A), IRAK1 (B), TRAF6 (C), IKKα (D), IKKβ (E), or p65 (F), along with an increasing amount of Vangl2 (0, 250, 500, and 1000 ng), was measured at 24 h after transfection and normalized to the Renilla luciferase activity.

(G and H) HEK293T cells were transfected with plasmids encoding HA-tagged Vangl2 and Flag-tagged key proteins in NF-κB signaling (Flag-IKKα, Flag-p65, Flag-TRAF6, Flag-IRAK1, Flag-MyD88 (G) and Flag-IKKβ (H), followed by immunoprecipitation with anti-Flag beads and immunoblot analysis with anti-HA. Throughout was the immunoblot analysis of whole-cell lysates (WCL) without immunoprecipitation.

(I) BMDMs were stimulated with LPS (100 ng/ml) for the indicated times. The cell lysates were subjected to immunoprecipitation with an anti-Vangl2 antibody or control IgG, followed by immunoblotting with the indicated antibodies.

(J) Interacting domains of Vangl2 and p65 predicted by ZDOCK server.

(K) Quantification of co-localization of p65 and Vangl2 in peritoneal macrophages.

(L) THP-1 were stimulated with LPS (100 ng/ml) for 8 h and the cells were isolated to cytoplasm and membrane fractions by kits. The cell lysates were subjected to immunoprecipitation with an anti-p65 antibody or control IgG, followed by immunoblotting with the indicated antibodies.

(M) HEK293T cells were transfected with Flag-tagged p65 and empty vector, Myc-tagged Vangl2 (FL) or Vangl2 truncation mutants. The cell lysates were subjected to immunoprecipitation with anti-Flag beads and immunoblotted with the indicated antibodies.

IP, immunoprecipitation; WCL, whole-cell lysate. Data are representative of three independent experiments.

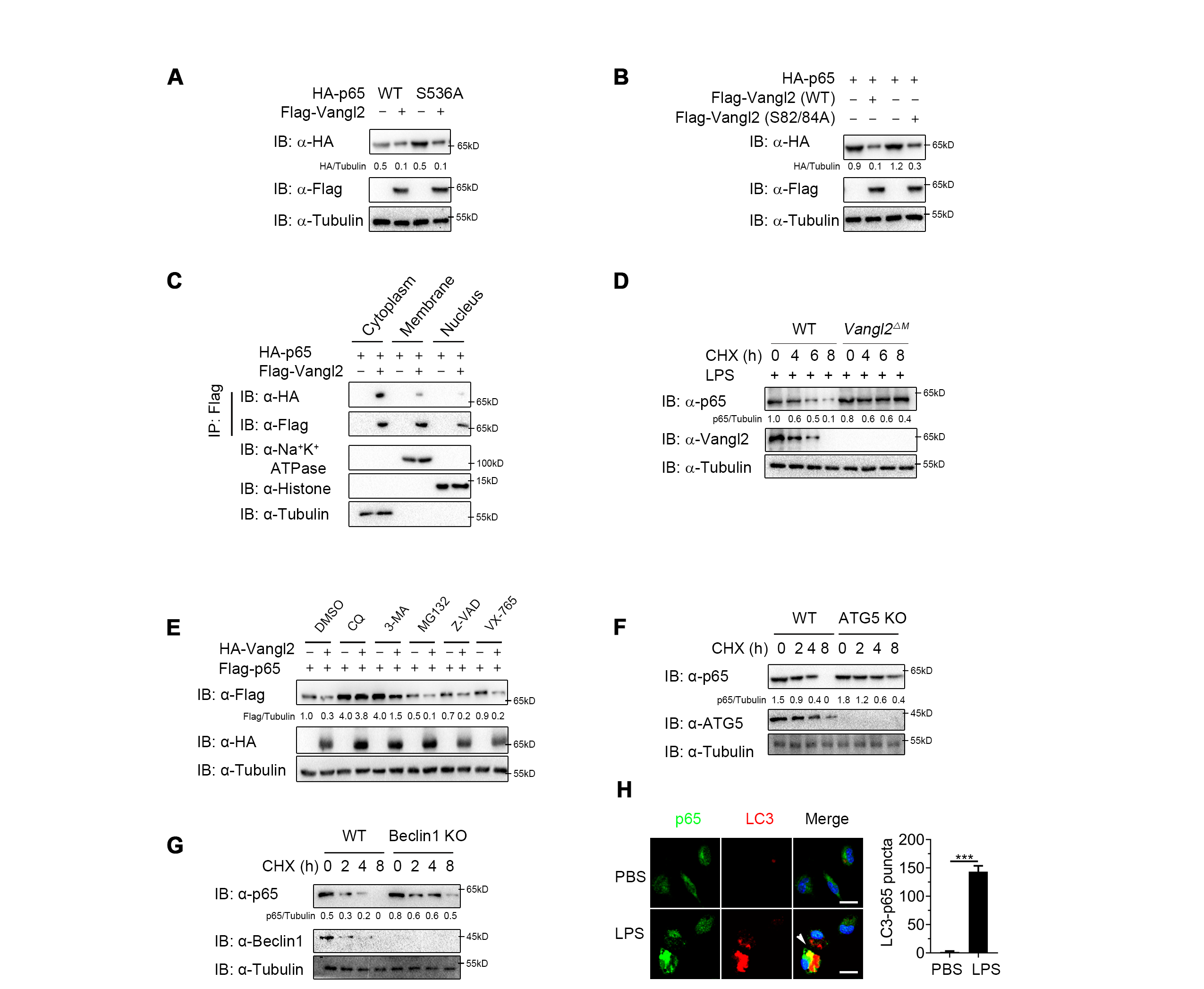

**Figure 4-figure supplement 4. Vangl2 promotes p65 degradation by autophagic pathway.**

(A) Immunoblot analysis of HEK293T cells transfected with HA-p65 (WT or S536A) and Flag-Vangl2.

(B) Immunoblot analysis of HEK293T cells transfected with HA-p65 and Flag-Vangl2 (S82/84A).

(C) HEK293T cells were transfected with Flag-tagged Vangl2 and empty vector, HA-tagged p65. The cells were isolated to cytoplasm, membrane and nucleus fractions by kits. Cell lysates were subjected to immunoprecipitation with anti-Flag beads and immunoblotted with the indicated antibodies.

(D) WT and Vangl2-deficient BMDMs were pretreated with LPS, then treated with CHX for the indicated times, and the expressions of p65 and Vangl2 were detected by immunoblot.

(E) HEK293T cells were transfected with Flag-p65, together with or without HA-Vangl2 plasmids, and treated with DMSO, MG132 (10 μM), CQ (50 μM), 3-MA (10 mM), VX-765 (10 mM) or Z-VAD (0.2 μM) for 6 h. The cell lysates were analyzed by immunoblot with indicated antibodies.

(F and G) WT and ATG5 KO (F) or Beclin1 KO (G) HEK293T cells were treated with CHX for the indicated times, and then the cell lysates were analyzed by immunoblot with indicated antibodies.

(H) Confocal microscopy of WT BMDMs treated with PBS and LPS. Statistics shown refer to the puncta formation by LC3-p65 in the indicated samples. Scale bar, 50 μm. (p65, green; LC3, red; DAPI, blue)

Data are representative of three independent experiments and are plotted as the mean ±SD. ****p*<0.001 vs. corresponding control.

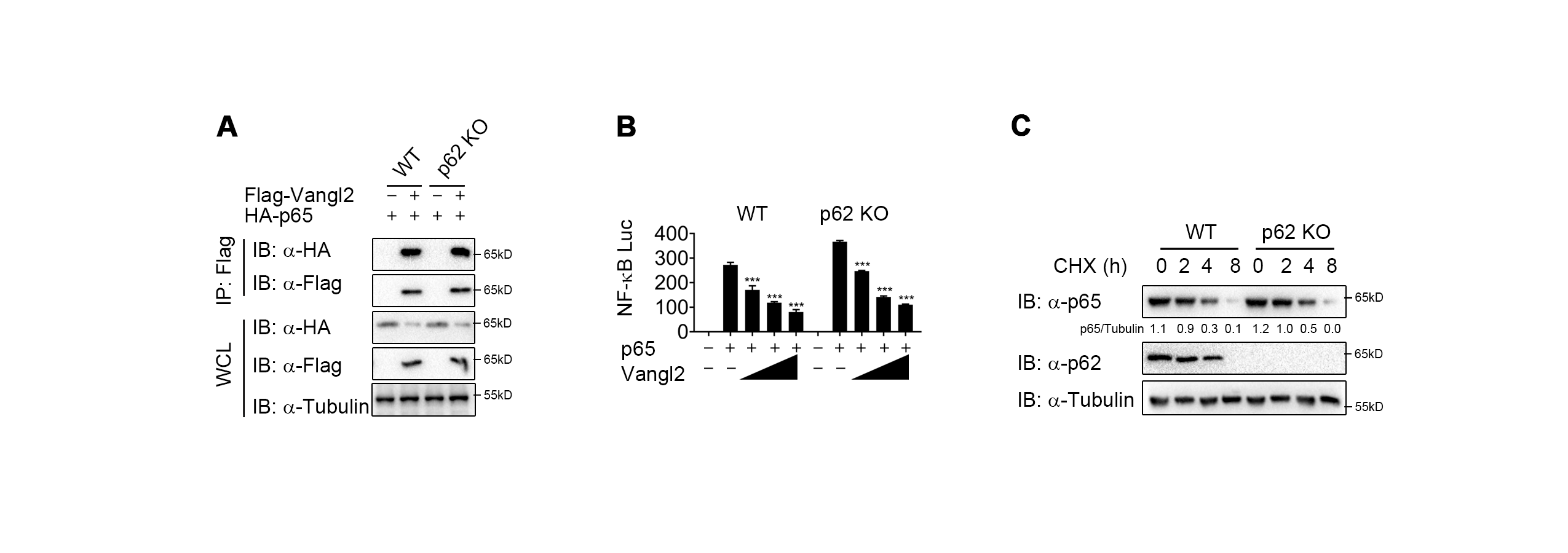

**Figure 5-figure supplement 5. Vangl2 promoted autophagic degradation of p65 is not mediated by cargo receptor p62.**

(A) WT and p62 KO HEK293T cells transfected with a vector expressing HA-p65 along with the empty vector or vector encoding Flag-Vangl2. The cell lysates were subjected to immunoprecipitation with anti-Flag beads and immunoblotted with the indicated antibodies.

(B) Luciferase activity in WT and p62 KO HEK293T cells transfected with plasmids encoding an NF-κB luciferase reporter and TK-Renilla reporter, together with p65 plasmid along with increasing amounts of Vangl2 plasmid, was measured at 24 h after transfection.

(C) WT and p62 KO HEK293T were treated with CHX for the indicated times. The cell lysates were immunoblotted with the indicated antibodies.

Data are representative of three independent experiments and are plotted as the mean ±SD. ****p*<0.001 vs. corresponding control.

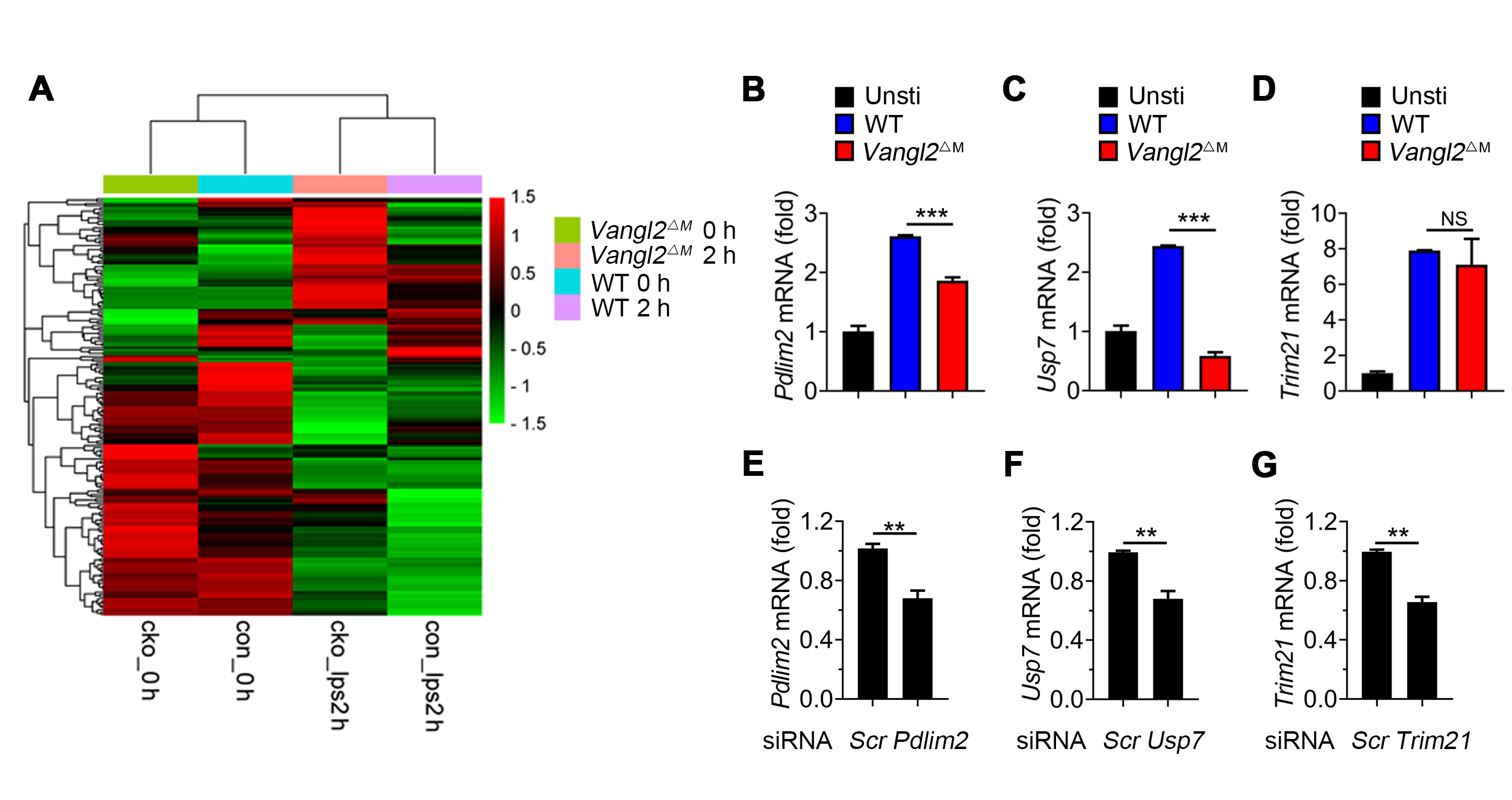

**Figure 7-figure supplement 6. Expression of candidate E3 ubiquitin ligases in WT and Vangl2-deficient BMDMs after LPS treatment.**

(A) Heatmap view of mRNA variations of E3 ubiquitin ligases in WT and Vangl2-deficient BMDMs treated with or without LPS.

(B-D) mRNA levels of *Pdlim2* (B), *Usp7* (C), and *Trim21* (D) in the spleens from LPS-treated WT and *Vangl2^△M^* mice were detected by qPCR (n≥4).

(E-G) mRNA levels of *Pdlim2* (E), *Usp7* (F), and *Trim21* (G) in HEK293T after transfection with *Pdlim2*, *Usp7*, and *Trim21* siRNA.

Data are representative of three independent experiments and are plotted as the mean ±SD. ***p*<0.01, ****p*<0.001 vs. corresponding control. NS, not significant.

**Supplementary Materials and Methods**

**Sepsis patients’ information**

Whole blood samples from patients who were diagnosed as sepsis induced by gram-negative bacterial infection were obtained from Nanfang Hospital, Southern Medical University. Criteria for enrolled patients followed by the international guidelines for management of sepsis and septic shock of 2016 (NFEC-2023-437).

| ID | Sex | Age | Admission diagnosis | CRP (mg/mL) | WBC (10^9^/L) | Heart rate (beats/min) |
| --- | --- | --- | --- | --- | --- | --- |
| 1# | F | 68 | COPD | 82.61 | 13.86 | 112 |
| 2# | F | 61 | Cholangitis | 42.58 | 23.99 | 121 |
| 3# | F | 72 | Femoral fracture | 60.31 | 18.23 | 127 |
| 4# | M | 57 | Diabetes | 55.32 | 14.33 | 122 |
| 5# | M | 68 | COPD | 62.61 | 13.99 | 124 |
| 6# | M | 72 | COPD | 52.58 | 28.23 | 140 |
| 7# | M | 78 | COPD | 70.32 | 24.33 | 128 |
| 8# | F | 82 | COPD | 45.32 | 15.34 | 110 |
| 9# | M | 25 | Health checkup | 10.23 | 6.55 | 65 |
| 10# | M | 28 | Health checkup | 5.84 | 7.56 | 68 |
| 11# | M | 27 | Health checkup | 15.32 | 5.57 | 72 |
| 12# | F | 29 | Health checkup | 14.21 | 6.85 | 75 |
| 13# | F | 26 | Health checkup | 8.22 | 4.56 | 67 |
| 14# | F | 27 | Health checkup | 5.82 | 6.02 | 66 |
| 15# | M | 30 | Health checkup | 12.32 | 5.81 | 73 |
| 16# | M | 31 | Health checkup | 9.81 | 5.73 | 77 |

**Immunoprecipitation and immunoblot analyses**

Cells were lysed by low-salt lysis buffer. For endogenous immunoprecipitation, whole-cell lysates were treated with indicated antibodies overnight and then incubated protein A/G beads (Pierce) for 4-6 h. For exogenous immunoprecipitation, whole-cell lysates were incubated with anti-FLAG or anti-Myc agarose gels. Immunoprecipitates were eluted with 2×SDS loading buffer after 5 times washing with low-salt lysis buffer. The proteins were dissolved in SDS loading buffer and boiled for 8-10 min. Then protein lysates resolved on SDS-PAGE gels and proteins were transferred to a polyvinylidene difluoride membrane (Millipore). After blocking with 5% (w/v) reagent-grade nonfat milk (Sigma), the membranes were incubated with the indicated antibodies (Table S1) overnight. For all blots, proteins were detected by EMD Millipore Luminata Western HRP Chemiluminescence Substrate.

**Flow cytometry analysis**

Mouse splenocytes were stained with indicated antibodies (Table S1) at 4 °C in RPMI containing 2% FBS for 30-60 min. All samples were detected by BD LSRFortessa flow cytometry analyzer (BD sciences). The data were analyzed via FlowJo X software (Tree Star).

**RNA Preparation and qPCR**

Total RNA was purified from stimulated cells and splenic tissue by the TRIzol reagent (Invitrogen), and cDNA was obtained using starscript II first-stand cDNA synthesis kit (GenStar, Beijing, China). Realtime PCR was performed on QuantStudio 6 flex (Thermo Fisher, Waltham, MA, USA) using RealStar green power mixture (GenStar, Beijing, China) with primers listed in table S2.

**Cellular fractionation**

Cells were collected by scraping, spun down and washed in pre-chilled PBS. For cytoplasmic and nuclear extracts were prepared by NE-PER Nuclear and Cytoplasmic Extraction Reagents (Thermo). Briefly, cytoplasmic were extracted by ice-cold CER I and CER II reagent, and nuclear were extracted by ice-cold CER I and CER II NER reagent (Tanaka, Grusby, & Kaisho, 2007). For cytosol and membrane, MELB buffer (20 mM 4-(2-Hydroxyethyl) piperazine-1-ethanesulfonic acid (HEPES) pH = 7.5, 100 mM sucrose, 2.5 mM MgCl2, 100 mM KCl) containing 0.025% digitonin was used to permeabilize cells to extract the cytosol fraction, and 1% digitonin buffer was used to extract cell membrane fraction (Liu et al., 2021).

**Table S1. Reagents and antibodies used in this study.**

| **Antibody Name** | **Source** | **Catalog number** |
| --- | --- | --- |
| Vangl2 | Santa Cruz | #sc-515187 |
| phosphor-IKK-α/β (Ser^178/180^) | Cell Signaling Technology | #2697 |
| IKK-α/β | Santa Cruz | #52932 |
| Pro-IL-1β | Cell Signaling Technology | #12507 |
| IL-1β (p17) | Cell Signaling Technology | #12242 |
| phosphor-p65 (Ser^536^) | Cell Signaling Technology | #3033 |
| p65 | Cell Signaling Technology | #8242 |
| Anti-mouse CD8a | eBioscience | 17-0081-82 |
| Beclin1 | proteintech | 11306-1-AP |
| Ndp52 | proteintech | 12229-1-AP |
| Atg5 | proteintech | 10181-2-AP |
| p62 | proteintech | 18420-1-AP |
| Anti-Flag | Sigma | A8592 |
| Anti-HA | Sigma | 12013819001 |
| Anti- Myc | Beijing Ray | RM1003 |
| Anti-mouse CD11b | eBioscience | 48-0112-82 |
| Anti-mouse F4/80 | eBioscience | 17-4801-82 |
| Anti-mouse Ly6C | eBioscience | 128003 |
| Anti-mouse Ly6G | eBioscience | 127603 |

**Table S2. Primers sequences for quantitative RT-PCR**

| **Description gene/protein** | **Forward Primer Sequence** | **Reverse Primer Sequence** |
| --- | --- | --- |
| Mouse-*Gapdh* | AGGTCGGTGTGAACGGATTTG | TGTAGACCATGTAGTTGAGGTCA |
| Mouse-*Vangl2* | TGAGGGCCTCTTCATCTCC | GCCCGTGGAGTTAATTGGT |
| Mouse-*Il1b* | CACAGCAGCACATCAACAAG | GTGCTCATGTCCTCATCCTG |
| Mouse-*Il6* | CCAGTTTGGTAGCATCCATC | CTCTGGGAAATCGTGGAAAT |
| Mouse-*Tnfa* | GACGTGGAACTGGCAGAAGAG | TTGGTGGTTTGTGAGTGTGAG |
| Mouse-*Pdlim2* | TGGGGCTTCCGAATTAGCG | CGCGTGTAGCATGTTCTCTG |
| Mouse-*Usp7* | TCGTCGCACATTGAGACGG | CTTGTCGGCATGGTTGGGAAT |
| Mouse-*Trim21* | GGGAGGAGGTCACCTGTTCTA | GGCACTCGGGACATGAACTG |
| Human-*Gapdh* | GGCTGTTGTCATACTTCTCATGG | CCCTATTCCCCACAACACAC |
| Human-*Vangl2* | AATCCCGAAAAGAAGGCTGT | CCCTATTCCCCACAACACAC |
| Human-*p65* | CCCACGAGCTTGTAGGAAAGG | GGATTCCCAGGTTCTGGAAAC |
| Human-*Pdlim2* | GCCCATCATGGTGACTAAGG | ATGGCCACGATTATGTCTCC |
| Human-*Usp7* | CCAGTGCAATGCTGAATCTGA | ACGACGACTGAACGACTTTTCAT |
| Human-*Trim21* | TCAGCAGCACGCTTGACAAT | GGCCACACTCGATGCTCAC |
